# Differential Interactions Between Human ACE2 and Spike RBD of SARS-CoV-2 Variants of Concern

**DOI:** 10.1101/2021.07.23.453598

**Authors:** Seonghan Kim, Yi Liu, Zewei Lei, Jeffrey Dicker, Yiwei Cao, X. Frank Zhang, Wonpil Im

## Abstract

Severe acute respiratory syndrome coronavirus 2 (SARS-CoV-2) is the causative agent of the current coronavirus disease 2019 (COVID-19) pandemic. It is known that the receptor-binding domain (RBD) of the spike protein of SARS-CoV-2 interacts with the human angiotensin-converting enzyme 2 (ACE2) receptor, initiating the entry of SARS-CoV-2. Since its emergence, a number of SARS-CoV-2 variants have been reported, and the variants that show high infectivity are classified as the variants of concern according to the US CDC. In this study, we performed both all-atom steered molecular dynamics (SMD) simulations and microscale thermophoresis (MST) experiments to characterize the binding interactions between ACE2 and RBD of all current variants of concern (Alpha, Beta, Gamma, and Delta) and two variants of interest (Epsilon and Kappa). We report that the RBD of the Alpha (N501Y) variant requires the highest amount of force initially to be detached from ACE2 due to the N501Y mutation in addition to the role of N90-glycan, followed by Beta/Gamma (K417N/T, E484K, and N501Y) or Delta (L452R and T478K) variant. Among all variants investigated in this work, the RBD of the Epsilon (L452R) variant is relatively easily detached from ACE2. Our results combined SMD simulations and MST experiments indicate what makes each variant more contagious in terms of RBD and ACE2 interactions. This study could help develop new drugs to inhibit SARS-CoV-2 entry effectively.

TOC Graphic

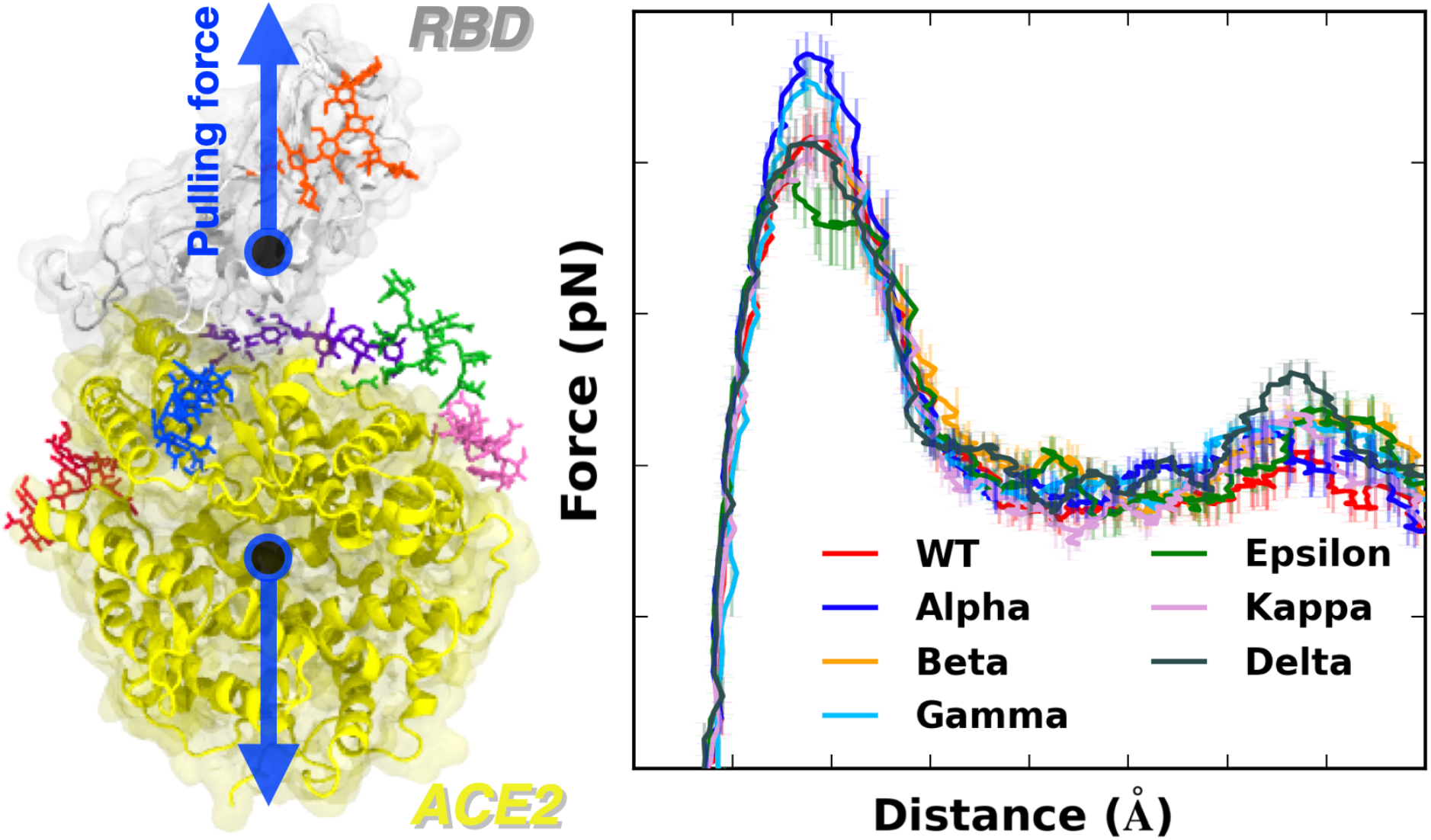

---

Reported in late 2019, severe acute respiratory syndrome coronavirus 2 (SARS-CoV-2) emerged and has rapidly infected people around the world. As of mid-July 2021, 192 million cases and 4.13 million deaths have been reported globally.^1^ Despite worldwide efforts to overcome the current coronavirus disease 2019 (COVID-19) pandemic, the rise of various SARS-CoV-2 variants may become futile the efficacy of vaccination and other countermeasures.

SARS-CoV-2 virus utilizes receptor-binding domain (RBD) of S1 protein, a part of trimeric spike (S) glycoprotein,^2–3^ for viral entry through the RBD interaction with the human receptor angiotensin-converting enzyme 2 (ACE2). Since ACE2 can interact with the RBD of both SARS-CoV-2 and SARS-CoV-1 (the virus caused the 2002-2004 SARS outbreak), there have been many studies not only to understand binding interactions between RBD and ACE2, but also to characterize the difference between SARS-CoV-1 and SARS-CoV-2.^4–6^

In September 2020, Alpha variant, lineage B.1.1.7, was first detected in southeast England and quickly became a populated lineage in the United Kingdom. The variant has been subsequently detected in the United States in December 2020.^7–8^ Beta variant, lineage B.1.351, was first detected in South Africa in May 2020 and found in the United States at the end of January 2021.^9^ At that time, there was another identified Gamma variant, which is known for lineage P.1,^10–11^ in the United States that was initially found in Japan from a traveler from Brazil. In November 2020, Epsilon variant, lineage B.1.427, was detected in California in the United States.^12^ Recently, two additional variants, Kappa (lineage B.1.617.1) and Delta (lineage B.1.617.2), first identified in India at the end of 2020, were detected in the United States.^13^ Since the emergence of diverse SARS-CoV-2 variants, Alpha, Beta, Gamma, and Delta variants are classified as the variants of concern by the US CDC due to their high infectivity.

To better understand the highly contagious characteristics of these variants, several studies have been performed experimentally and computationally.^14–16^ For example, Tian et al. conducted an experimental and computational study to capture the role of N501Y mutation in Alpha, Beta, and Gamma variants.^14^ They suggested that the π-π interactions and π-cation interactions are responsible for the enhanced interactions between RBD and ACE2. However, only the N501Y mutation was examined in their study, although other potentially important mutations have emerged. More recently, Socher et al. performed energy decomposition analysis from molecular dynamics simulations to compare the interaction energies between ACE2 and RBD of Alpha, Beta, and Gamma variants.^15^ They investigated each specific mutation, N501Y, K417N/T, and E484K, and reported that F486, Q498, T500, and Y505 in RBD are important residues across viral variants in the RBD-ACE2 interface.

In this study, using all-atom steered molecular dynamics (SMD) simulations and microscale thermophoresis (MST) experiments (see Supporting Information Methods), we report the differential interactions between human ACE2 and RBD of SARS-CoV-2 of all variants of concern (Alpha, Beta, Gamma, and Delta) as well as two variants of interest (Epsilon and Kappa). The study also provides a better understanding of such differences at the molecular level.

To gain molecular insight into the difference of all variants that are classified as the variants of concern (Alpha (first identified in United Kingdom, B.1.1.7: N501Y), Beta (first identified in South Africa, B.1.351: K417N, E484K, N501Y), Gamma (first identified in Japan/Brazil, P.1: K417T, E484K, N501Y), and Delta (first identified in India, B.1.617.2: L452R, T478K)) and two additional variants of interest (Epsilon (first identified in US-California, B.1.427: L452R) and Kappa (first identified in India, B.1.617.1: L452R, E484Q)), pulling force analysis was performed on each RBD-ACE2 complex (**Figure 1A**) as a function of distance (*D*) between the centers of mass of RBD and ACE2 proteins. Our fully-glycosylated S RBD-ACE2 complex model (**Figure 1B,C**) was employed for the pulling simulation.^17^ As shown in **Figure 1A**, most variants have increased force profiles than WT except for the Epsilon variant, indicating that the variants have strengthened interactions with ACE2. It should be noted that the amount of initial force at *D* = 53 Å shows a good match with our previous WT study,^4^ where we utilized only the N-linked glycan (N-glycan) core structure for all N-glycans. In this study, we used the most probable N-glycan structures (**Figure 1C**) that are larger than the core structure.

**Figure 1.**
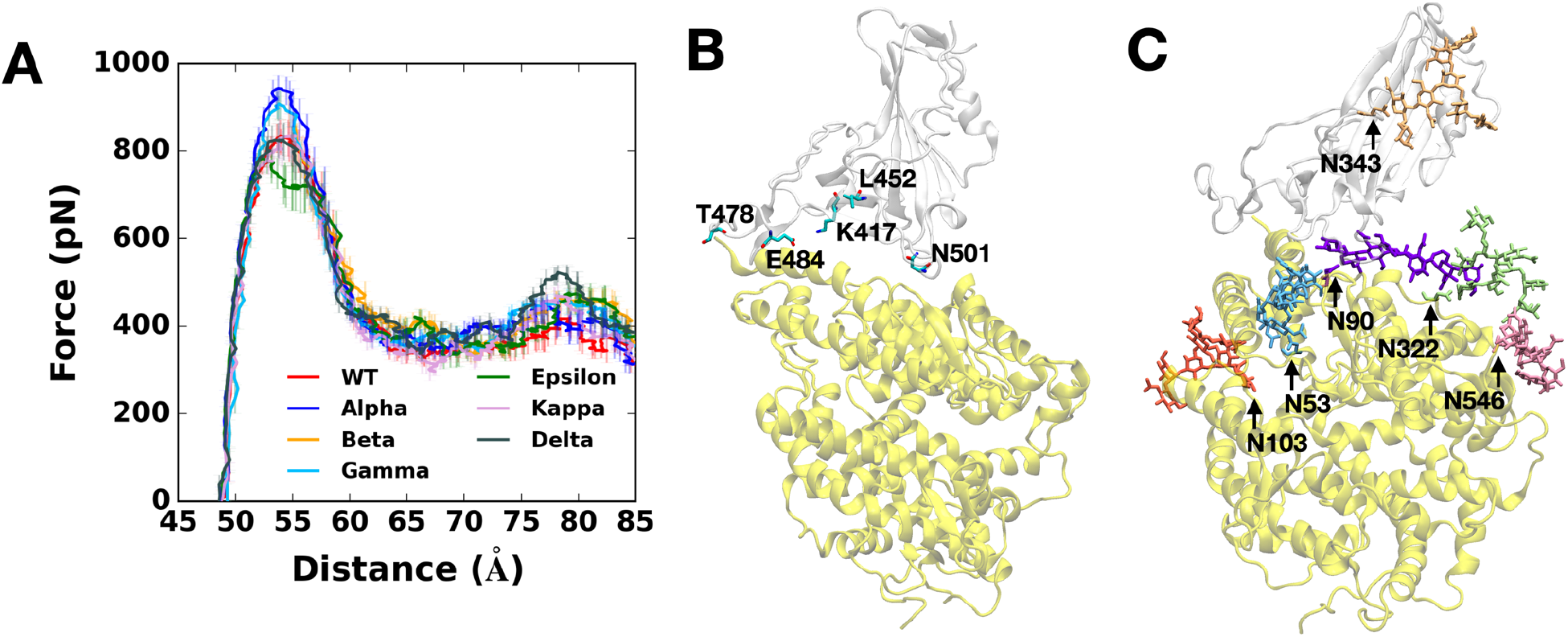
(A) Average force profiles of WT (red), Alpha (blue), Beta (orange), Gamma (sky blue), Epsilon (green), Kappa (pink), and Delta (gray) variants as a function of the distance between the centers of mass of RBD and ACE2. (B) Initial snapshot of WT. Residues subjected to each mutation are shown as solid sticks (N501, K417, E484, L452, and T478). RBD and ACE2 are respectively colored in light gray and yellow. All N-glycans, water, and ions are hidden for clarity. (C) Initial snapshot of WT with clockwise 90° rotation along the normal from (B). All N-glycans are depicted in different colors. Any other residues, water, and ions are not shown for clarity.

**Figure 1A** shows that, at *D* = 53 Å, Alpha variant clearly requires the highest initial force to pull the RBD-ACE2 complex in the opposite direction. The difference can be explained in **Figure 2B**, a two-dimensional contact map between RBD^Alpha^ and ACE2 at *D* = 53 Å, where RBD Y501 presents increased interactions with ACE2 Q42, Y41, and D38. Such contacts are decreased or even lost in the case of RBD^WT^ or RBD^Epsilon^ lacking the N501Y mutation (**Figure 2A,D**). To quantify the contact frequency between RBD residue 501 (N501 for WT, Epsilon, Kappa, and Delta; Y501 for Alpha, Beta, and Gamma) and ACE2, the number of heavy atom contacts was calculated (**Figure 3A**). The contact was counted if RBD residue 501 is positioned within 4.5 Å of heavy atoms of key interacting residues of ACE2 protein. Notably, Y501 of Alpha, Beta, and Gamma variants retain more contacts (about 40%) than N501 of WT, Epsilon, Kappa, and Delta variants. As shown in **Figure 3B,C**, Alpha Y501 is located closer to ACE2 Y41 and K353 than WT N501 at *D* = 53 Å, and thus, it has the π-π and π-cation interactions with neighboring Y41 and K353. On top of the Y501-ACE2 interactions, RBD^Alpha^ also contains the highest amounts of contacts with ACE2 N90-glycan (**Figure S3**), which could be the reason why it has been reported as the most common lineages by June 19, 2021, among the estimated proportions of SARS-CoV2 lineages according to the US CDC,^18^ even though this study considers only single RBD out of trimeric SARS-CoV2 S protein.

**Figure 2.**
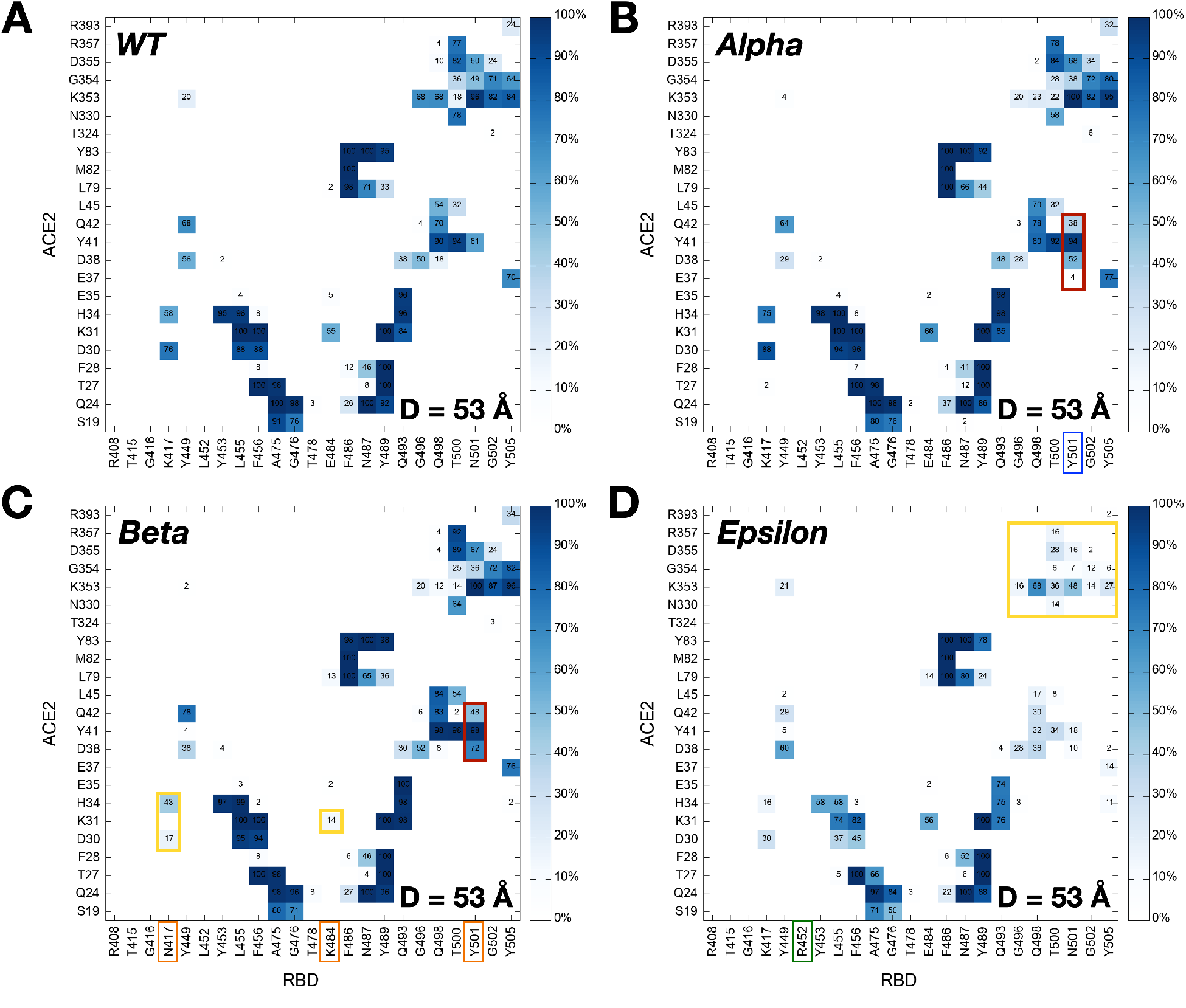
Two-dimensional contact maps at *D* = 53 Å. (A) Interacting residue pairs between RBD^WT^ and ACE2. RBD residues subjected to mutation are shown in colored boxes at the bottom: (B) blue for Alpha, (C) orange for Beta, and (D) green for Epsilon. The contact frequency is numbered with colors from light blue to dark blue. Dark red and yellow colors on the map respectively represent increased and decreased interactions between RBD and ACE2 upon mutations.

**Figure 3.**
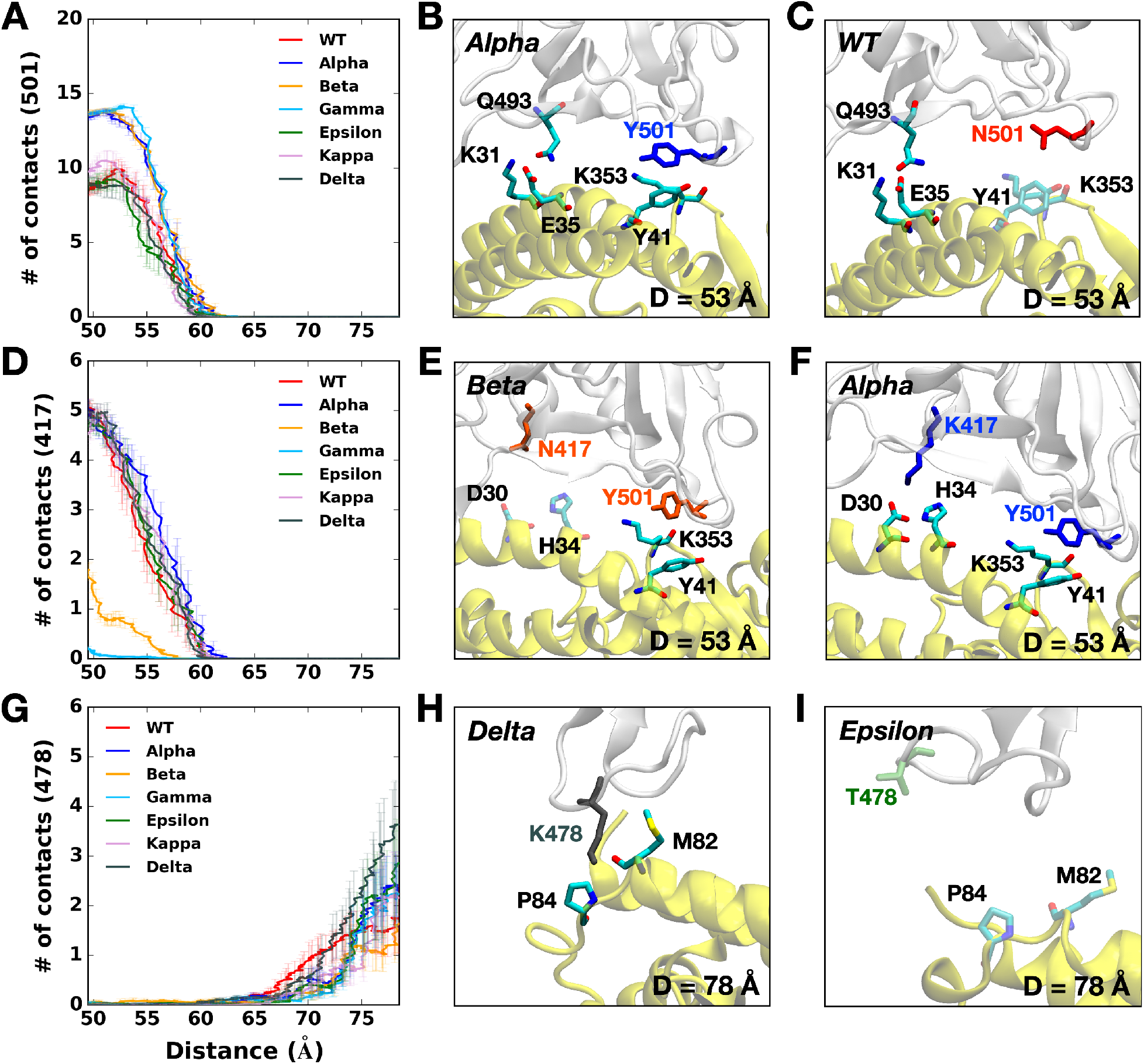
(A) The average number of contacts between RBD residue 501 and ACE2. (B and C) Representative snapshots at *D* = 53 Å of (B) Alpha variant and (C) WT. (D) The average number of contacts between RBD residue 417 and ACE2 and (E and F) their interacting residue pairs at *D* = 53 Å of (E) Beta and (F) Alpha variants. (G) The average number of contacts between RBD residue 478 and ACE2 and (H and I) key interaction pairs at *D* = 78 Å of (H) Delta and (I) Epsilon variants. The overall color scheme is the same as in **Figure 1**, and each mutated residue in each variant is shown using the same colors (i.e., red for WT, blue for Alpha, orange for Beta, green for Epsilon, and gray for Delta). Interacting residues are depicted as the solid sticks and residues losing their interactions are shown as the transparent sticks. RBD and ACE2 are presented in light gray and yellow, respectively.

The force profiles of Beta and Gamma variants at *D* = 53 Å present weaker maximum forces than the Alpha variant, although they show higher forces than WT at the same distance (**Figure 1A**). As shown in **Figure 2B,C**, Alpha and Beta variants include N501Y mutation, while Beta variant involves two additional mutations, K417N and E484K. Clearly, compared to WT or Epsilon, Y501 of Beta variant has increased interactions (colored in dark red box) with ACE2 D38, Y41, and Q42, similar to Alpha variant. However, it entails decreased contact frequency (shown as the yellow box) between RBD^Beta^ N417 and ACE2 D30/H34, as well as RBD^Beta^ K484 and ACE2 K31, which could be explained why Beta has relatively weaker interactions than Alpha. Gamma variant also shows decreased contact numbers similar to Beta due to its K417T mutation (**Figure S2A**). The only difference between Gamma and Beta is the K417 mutation, i.e., K417T vs. K417N. **Figure 3D** compares the number of contacts of residue 417 of all variants that are in contact with heavy atoms of key interacting residues of ACE2. While all other variants containing K417 (i.e., WT, Alpha, Epsilon, Kappa, and Delta) display some contacts between RBD and ACE2 from 50 to 60 Å, few interactions were found for the Beta variant. The sidechain-shortening mutation from lysine to asparagine could have an impact on the RBD-ACE2 interface, resulting in fewer interactions at the same distance (**Figure 3E,F**). Interestingly, T417 of Gamma shows almost no interaction because threonine is even shorter than N417 of Beta. The weakened interactions of RBD^Beta^ N417 and RBD^Gamma^ T417 could make them less contagious than the Alpha variant, while the N501Y mutation still allows them to have a strong enough potential to interact with ACE2. This could explain why/how the Gamma variant took the second-highest portion by June 5, 2021, among the estimated proportions of SARS-CoV2 lineages, provided by the US CDC.^18^

Although most variants show similar maximum forces around *D* = 53 Å, the Epsilon variant shows decreased forces with more fluctuations than other variants (**Figure 1A**). The two-dimensional contact map in **Figure 2D** confirms its distinct interactions at *D* = 53 Å, as it shows the least number of contacts between RBD^Epsilon^ and ACE2 (n.b., the yellow box represents deceased interactions). For example, K353 residue of all other variants is actively interacting with ACE2 Q493, Q496, Q498, T500, N/Y501, G502, and Y505 (**Figure 2A,B,C** and **Figure S2A,B,C**). K353 of Epsilon, however, lost its contact with corresponding residues at least by 50%. To investigate the mechanism behind such a big difference, the contact analysis in between RBD residues was performed, where the influence of the L452R mutation was examined by checking its contacts with surrounding residues, L450 and L492 (**Figure S4**). Interestingly, mutated R452 interacts more with L450 (**Figure S4C**) and less with L492 (**Figure S4A**) simultaneously. Note that L450 and L492 are positioned in different β-strands (**Figure S4B,D** colored in green and orange, respectively), and the L452R mutation makes the RBD-ACE2 interface unstable by shortening each β-strand (i.e., the length of interacting β-strands of Epsilon variant is decreased by almost half). Because of such a less stable RBD structure, the Epsilon variant appears to be detached from ACE2 easier than WT. Indeed, K353 of Epsilon variant lost contacts with ACE2 Q498 and Y505 at *D* = 55 Å (**Figure S4D**), but WT holds their interactions at the same distance (**Figure S4B**). As of June 29, 2021, according to the US CDC, the Epsilon variant deescalated from the variants of concern and became the variants of interest since its considerable decrease in terms of lineage proportion in the United States.

Newly reported Kappa and Delta variants display the same L452R mutation as Epsilon, but each variant contains an additional mutation, E484Q (Kappa) or T478K (Delta). Even though both Epsilon and Kappa/Delta variants share L452R mutation, Kappa and Delta variants show similar trends in force profiles to WT from *D* = 55 Å to 75 Å (**Figure 1A**). Specifically, the two-dimensional contact maps of Kappa and Delta variants (**Figure S2B,C**) display almost identical interaction patterns to WT between ACE2 K353 and RBD residues (i.e., Q493, Q496, Q498, T500, N/Y501, G502, and Y505), while Epsilon variant at *D* = 53 Å (**Figure 2A,D**) does not. It should be noted that this difference between Kappa/Delta and Epsilon might stem from the limitation in our model in this study, as we only employed L452R mutation in RBD for Epsilon variant without D614G mutation.

The Delta variant, interestingly, shows distinct features that are not found in other variants. Upon the T478K mutation, it requires the highest force for the RBD-ACE2 complex to be completely dissociated at *D* = 78 Å (**Figure 1A**). In order to see what makes the difference, the number of contacts between RBD residue 478 and heavy atoms of selected key interacting residues of ACE2 was calculated. As shown in **Figure 3G**, RBD^Delta^ exclusively makes more contacts with ACE2 than other variants. **Figure 3H** shows that Delta K478 retains contacts with ACE2 P84 and M82 at *D* = 78 Å, but Epsilon T478 already lost such interactions. It is possible that residue 478 located in the flexible loop could first have a chance to contact with ACE2, and the stronger interactions of Delta K478 with ACE2 could explain the reason why the proportion of Delta variant is recently dramatically being increased with high infectivity. The Delta variant recently became the current variants of concern, and it took the highest portion among the estimated variant proportions as of July 3, 2021, according to the US CDC.^18^

To validate the SMD simulation results, we conducted an experimental protein binding assay using microscale thermophoresis (MST). MST detects molecular binding kinetics based on the thermophoretic movement of molecules induced by a microscopic temperature gradient inside a glass capillary generated by an infrared laser.^19^ MST has been used for detecting viral protein-receptor interactions,^20^ including SARS-CoV-2 S proteins.^21^ In our assay, human recombinant ACE2 was fluorescently labeled, and various RBD variants were titrated in a two-fold fashion and mixed with the ACE2. The MST signal was first converted to saturated fraction data and subsequently fitted to a first-order 1:1 binding kinetics model using the manufacturer’s software (**Figure S5**). The binding affinities of ACE2 and RBD^WT^ were detected to be 27.5 ± 4.8 nM (**Figure 4**). This value is in agreement with a reported Kd range of 5-40 nM measured by surface plasmon resonance.^22^ Importantly, our MST data indicate that Alpha variant binds ACE2 with a 2.3-fold higher affinity (11.8 ± 0.8 nM) than WT. The rest of the variants show slightly different affinity from WT. Beta and Delta variants display approximately 20-30% higher affinities than WT, and Epsilon variant shows a 15% lower affinity than WT. In **Figure 4**, Kd from MST experiments were directly compared with the FWT/F ratio from the SMD simulations, where FWT and F are the maximum pulling forces of WT and each variant around D = 53 Å (**Figure 1A**). Our MST affinity data are consistent with the SMD simulation data, indicating Alpha and Epsilon variants possess the strongest and weakest binding to ACE2, respectively.

**Figure 4.**
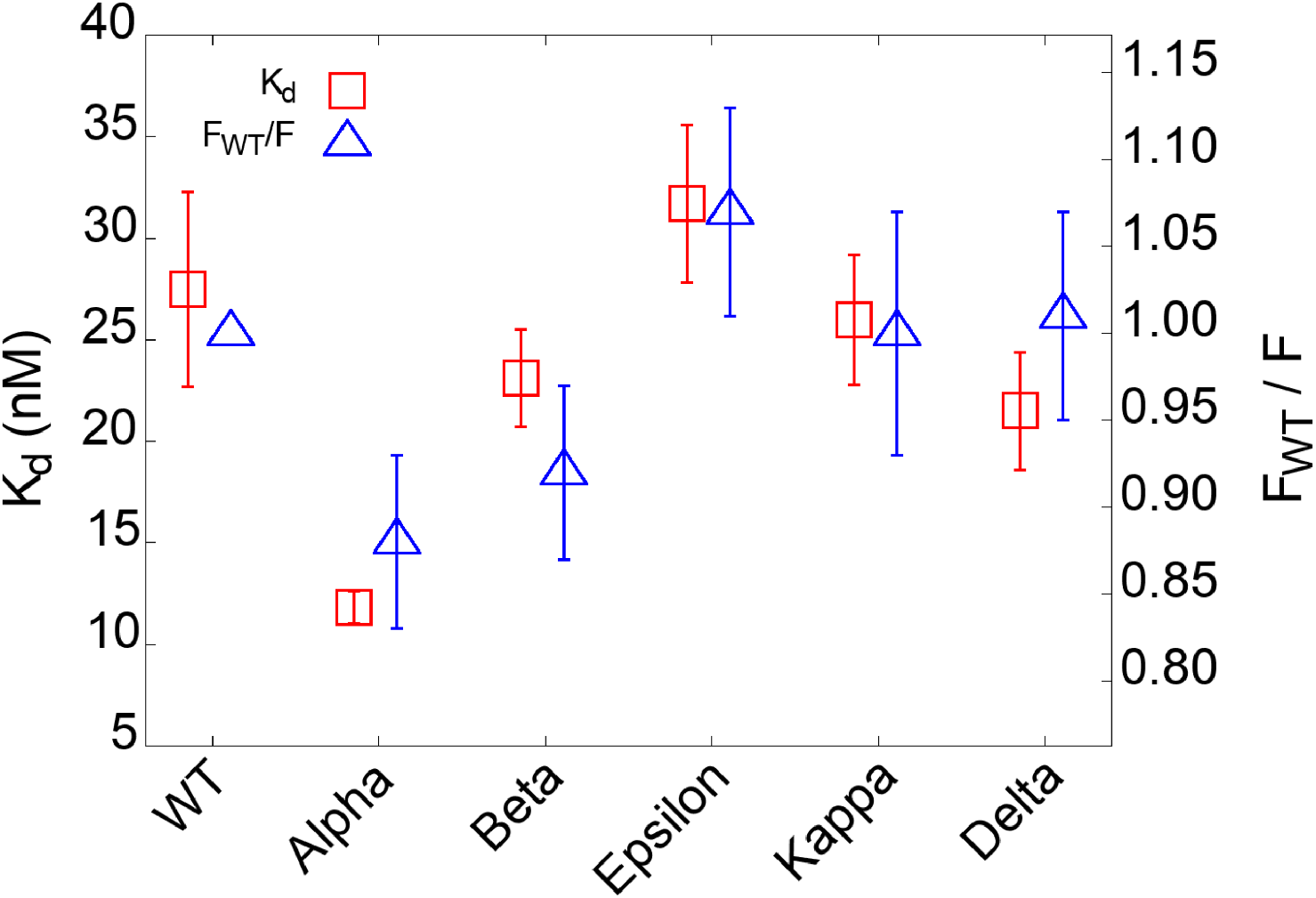
Binding affinities between RBD variants and ACE2 and its comparison with the simulation results. K_d_ is obtained from microscale thermophoresis experiments. F_WT_/F is a ratio, where FWT and F are the respective maximum pulling force of WT and of each variant obtained from the SMD simulations.

In summary, we characterized interactions between ACE2 and RBD of all variants that the US CDC classifies as the variants of concern and variants of interest. Our results indicate that Alpha variant requires the highest force for initial separation from ACE2, followed by Beta and Gamma variants or Delta variant. K417N/T mutations of Beta and Gamma appear to make the RBD-ACE2 interactions less strong compared to Alpha variant. In addition, Epsilon variant is likely to be relatively easily dissociated from ACE2 than others due to its destabilized RBD structure upon the L452R mutation. In addition, Delta variant specifically shows stronger interactions with ACE2 than other variants at a relatively far distance between RBD and ACE2. Our MST experiments show consistent results with the simulation results, where Alpha and Epsilon variants display the strongest and weakest binding to ACE2, respectively. This study provides valuable information that distinguishes important features of all variants and their interactions with ACE2.

## ACKNOWLEDGEMENTS

This work was supported in part by NIH R15AI133634 and NSF 1804117 (to X.F.Z.), NIH R21 AI163708 (to X.F.Z. and W.I.), NIH GM138472, and MCB-1810695 (to W.I.), and an internal grant from Lehigh University (to X.F.Z, and W.I.).

## REFERENCES

1 World Health Organization. WHO Coronavirus (COVID-19) Dashboard (accessed on July 23, 2021). https://covid19.who.int.

2 Shang, J.; Ye, G.; Shi, K.; Wan, Y.; Luo, C.; Aihara, H.; Geng, Q.; Auerbach, A.; Li, F. Structural basis of receptor recognition by SARS-CoV-2. Nature 2020, 581 (7807), 221–224.

3 Wrapp, D.; Wang, N.; Corbett, K. S.; Goldsmith, J. A.; Hsieh, C. L.; Abiona, O.; Graham, B. S.; McLellan, J. S. Cryo-EM structure of the 2019-nCoV spike in the prefusion conformation. Science 2020, 367 (6483), 1260–1263.

4 Cao, W.; Dong, C.; Kim, S.; Hou, D.; Tai, W.; Du, L.; Im, W.; Zhang, X. F. Biomechanical characterization of SARS-CoV-2 spike RBD and human ACE2 protein-protein interaction. Biophys. J. 2021, 120 (6), 1011–1019.

5 Hoffmann, M.; Kleine-Weber, H.; Schroeder, S.; Krüger, N.; Herrler, T.; Erichsen, S.; Schiergens, T. S.; Herrler, G.; Wu, N. H.; Nitsche, A.; Müller, M. A.; Drosten, C.; Pöhlmann, S. SARS-CoV-2 Cell Entry Depends on ACE2 and TMPRSS2 and Is Blocked by a Clinically Proven Protease Inhibitor. Cell 2020, 181 (2), 271–280.e8.

6 Wang, Q.; Zhang, Y.; Wu, L.; Niu, S.; Song, C.; Zhang, Z.; Lu, G.; Qiao, C.; Hu, Y.; Yuen, K. Y.; Wang, Q.; Zhou, H.; Yan, J.; Qi, J. Structural and Functional Basis of SARS-CoV-2 Entry by Using Human ACE2. Cell 2020, 181 (4), 894–904.e9.

7 Tang, J. W.; Tambyah, P. A.; Hui, D. S. C. Emergence of a new SARS-CoV-2 variant in the UK. J. Infect. 2021, 82 (4), e27–e28.

8 Kirby, T. New variant of SARS-CoV-2 in UK causes surge of COVID-19. Lancet Respir. Med. 2021, 9 (2), e20–e21.

9 Tegally, H.; Wilkinson, E.; Giovanetti, M.; Iranzadeh, A.; Fonseca, V.; Giandhari, J.; Doolabh, D.; Pillay, S.; San, E. J.; Msomi, N.; Mlisana, K.; von Gottberg, A.; Walaza, S.; Allam, M.; Ismail, A.; Mohale, T.; Glass, A. J.; Engelbrecht, S.; Van Zyl, G.; Preiser, W.; Petruccione, F.; Sigal, A.; Hardie, D.; Marais, G.; Hsiao, N.-y.; Korsman, S.; Davies, M.-A.; Tyers, L.; Mudau, I.; York, D.; Maslo, C.; Goedhals, D.; Abrahams, S.; Laguda-Akingba, O.; Alisoltani-Dehkordi, A.; Godzik, A.; Wibmer, C. K.; Sewell, B. T.; Lourenço, J.; Alcantara, L. C. J.; Kosakovsky Pond, S. L.; Weaver, S.; Martin, D.; Lessells, R. J.; Bhiman, J. N.; Williamson, C.; de Oliveira, T. Detection of a SARS-CoV-2 variant of concern in South Africa. Nature 2021, 592 (7854), 438–443.

10 Faria, N. R.; Claro, I. M.; Candido, D.; Franco, L. M.; Andrade, P. S.; Coletti, T. M.; Silva, C. A.; Sales, F. C.; Manuli, E. R.; Aguiar, R. S. Genomic characterisation of an emergent SARS-CoV-2 lineage in Manaus. Science 2021, 372 (6544), 815–821.

11 Voloch, C. M.; da Silva Francisco, R., Jr.; de Almeida, L. G. P.; Cardoso, C. C.; Brustolini, O. J.; Gerber, A. L.; Guimarães, A. P. C.; Mariani, D.; da Costa, R. M.; Ferreira, O. C., Jr.; Frauches, T. S.; de Mello, C. M. B.; Leitão, I. C.; Galliez, R. M.; Faffe, D. S.; Castiñeiras, T.; Tanuri, A.; de Vasconcelos, A. T. R. Genomic characterization of a novel SARS-CoV-2 lineage from Rio de Janeiro, Brazil. J. Virol. 2021, 95 (10), e00119–21.

12 Zhang, W.; Davis, B. D.; Chen, S. S.; Sincuir Martinez, J. M.; Plummer, J. T.; Vail, E. Emergence of a Novel SARS-CoV-2 Variant in Southern California. JAMA. 2021, 325 (13), 1324–1326.

13 Singh, J.; Rahman, S. A.; Ehtesham, N. Z.; Hira, S.; Hasnain, S. E. SARS-CoV-2 variants of concern are emerging in India. Nat. Med. 2021, 27, 1131–1133.

14 Tian, F.; Tong, B.; Sun, L.; Shi, S.; Zheng, B.; Wang, Z.; Dong, X.; Zheng, P. Mutation N501Y in RBD of Spike Protein Strengthens the Interaction between COVID-19 and its Receptor ACE2. bioRxiv 2021.

15 Socher, E.; Conrad, M.; Heger, L.; Paulsen, F.; Sticht, H.; Zunke, F.; Arnold, P. Decomposition of the SARS-CoV-2-ACE2 interface reveals a common trend among emerging viral variants. bioRxiv 2021.

16 Harvey, W. T.; Carabelli, A. M.; Jackson, B.; Gupta, R. K.; Thomson, E. C.; Harrison, E. M.; Ludden, C.; Reeve, R.; Rambaut, A.; Peacock, S. J.; Robertson, D. L.; Consortium, C.-G. U. SARS-CoV-2 variants, spike mutations and immune escape. Nat. Rev. Microbiol. 2021, 19 (7), 409–424.

17 Woo, H.; Park, S.-J.; Choi, Y. K.; Park, T.; Tanveer, M.; Cao, Y.; Kern, N. R.; Lee, J.; Yeom, M. S.; Croll, T. I.; Seok, C.; Im, W. Developing a Fully Glycosylated Full-Length SARS-CoV-2 Spike Protein Model in a Viral Membrane. J. Phys. Chem. B 2020, 124 (33), 7128–7137.

18 Centers for Disease Control and Prevention. Estimated Proportions of SARS-CoV-2 Lineages (accessed on July 23, 2021). https://covid.cdc.gov/covid-data-tracker/#variant-proportions.

19 Plach, M. G.; Grasser, K.; Schubert, T. MicroScale Thermophoresis as a tool to study protein-peptide interactions in the context of large eukaryotic protein complexes. Bio-Protoc. 2017, 7 (23), e2632–e2632.

20 Walls, A.; Tortorici, M. A.; Bosch, B. J.; Frenz, B.; Rottier, P. J.; DiMaio, F.; Rey, F. A.; Veesler, D. Crucial steps in the structure determination of a coronavirus spike glycoprotein using cryo-electron microscopy. Protein Sci. 2017, 26 (1), 113–121.

21 Petruk, G.; Puthia, M.; Petrlova, J.; Samsudin, F.; Strömdahl, A.-C.; Cerps, S.; Uller, L.; Kjellström, S.; Bond, P. J.; Schmidtchen; Artur. SARS-CoV-2 Spike protein binds to bacterial lipopolysaccharide and boosts proinflammatory activity. J. Mol. Cell. Biol. 2020, 12 (12), 916–932.

22 Liu, H.; Zhang, Q.; Wei, P.; Chen, Z.; Aviszus, K.; Yang, J.; Downing, W.; Jiang, C.; Liang, B.; Reynoso, L. The basis of a more contagious 501Y. V1 variant of SARS-COV-2. Cell Res. 2021, 31, 720–722.

