## Supporting Information for "Differential Interactions Between Human ACE2 and Spike RBD of SARS-CoV-2 Variants of Concern"

### Computational Methods

A fully-glycosylated SARS-CoV-2 RBD and ACE2 complex was obtained from COVID-19 Protein Library in CHARMM-GUI Archive (6vsb_1_1_1_6vw1.pdb).[1] The complex includes 6 N-linked glycans: five glycans in ACE2 (Asn53, Asn90, Asn103, Asn322, and Asn546) and one glycan in RBD (Asn343). For system generation, parameter setup, and corresponding mutations, we utilized CHARMM-GUI *Solution Builder*.[2-3] From the WT RBD structure, each variant was modeled with the following mutations: Alpha (N501Y), Beta (K417N, E484K, N501Y), Gamma (K417T, E484K, N501Y), Epsilon (L452R), Kappa (L452R, E484Q), and Delta (L452R, T478K). The CHARMM36(m) force field[4-5] for protein and carbohydrates with TIP3P water model[6] was used with 0.15 M of $K^+$ and $Cl^-$ ions for mimicking physiological conditions. The system size was determined to be large enough (about 190 Å ×190 Å × 190 Å) to have the proteins solvated enough when they are fully unattached. The total number of atoms is approximately 550,000.

The overall simulation details are nearly identical to our previous work.[7] NAMD simulation software[8] was used for the pulling simulations with the COLVARS method. As an initial condition, the SARS-CoV-2 RBD and ACE2 complex structures were aligned along the X-axis, and the center of mass (COM) of each protein was calculated to apply the external force on the proteins. The effective force acting on the COMs of both proteins can be calculated through the following equation:

$$U(\mathbf{r}_1, \mathbf{r}_2, \mathbf{r}_3, \ldots, t) = \frac{1}{2}k[vt - \mathbf{R}(t) \cdot \mathbf{n}]^2$$

where $k$ is the spring constant, $v$ is the moving speed of the spring potentials (also called dummy atoms), $\mathbf{R}(t)$ is the current position of the selected protein COM, and $\boldsymbol{n}$ is the COM-COM unit vector. This force enables the spring-connected proteins to move in the opposite directions to pull away two proteins. The moving speed of proteins was set to 0.5 Å/ns along the X-axis, and a spring constant of 5 kcal/mol/Å$^2$ was applied to the COM of each protein to have both proteins move along the X direction and restrict moving along the Y and Z directions. For better statistics, 20 independent simulations for each system were performed (140 systems total, 20 replicas of 7 variants) with at least 40 ns of each simulation run. The pulling simulations stopped when the RBD and ACE2 were completely detached from each other.

The van der Waals interactions were switched off smoothly over 10-12 Å using a force-based switching function.[9] The electrostatic interactions were calculated by the particle-mesh Ewald method with a mesh size of 1 Å.[10] To constrain bond lengths involving hydrogen atoms, the SHAKE algorithm was used.[11] The simulation time-step was set to 4 fs with the hydrogen mass repartitioning method.[12-13] Equilibration simulations were performed with the NVT (constant particle number, volume, and temperature) ensemble where positional and dihedral restraints were employed. The restraint was gradually decreased during the equilibration simulations. The NPT (constant particle number, pressure, and temperature) ensemble was then applied for the production runs, where the Langevin piston method[14] was used for the pressure control. The entire simulation temperature was set to 303.15 K with the Langevin damping control method.

### Experimental Methods

Recombinant human ACE2 protein (GenBank accession: AF291820.1, Sino Biological 10108-H08H; Wayne, PA) was labeled with RED-NHS (2nd Generation) dye using the Monolith Protein

Labeling Kit (NanoTemper Technologies, MO-L011, München, Germany). Labeled ACE2 (5 nM, final concentration) was mixed with the RBD proteins (WT or variants, 2-fold diluted in a 15-step starting from 1.5-4 μM) in PBS buffer supplanted with 0.1 % Pluronic® F-127. The RBD proteins include: WT (ACRObiosystems, SPD-C52H3, Newark, DE, GenBank accession: QHD43416.1), Alpha (N501Y, ACRObiosystems, SPD-C52Hn), Beta (K417N, E484K, N501Y, ACRObiosystems, SPD-C52Hp), Epsilon (L452R, Sino Biological, 40592-V08H28), Kappa (L452R, E484Q, Sino Biological, 40592-V08H88), and Delta (L452R, T478K, Sino Biological, 40592-V08H90). The mixed RBD+ACE2 samples were separately loaded into 16 premium glass capillaries (NanoTemper Technologies, MO-K025). The 16 capillaries were then placed in the reaction chamber in the order of concentration. Microscale thermophoresis (MST) measurements were conducted on a Monolith NT.115 instrument (NanoTemper Technologies) at 20% excitation power at 24 °C. The measurement was repeated at least three times. $K_d$ calculations were performed using the MO Affinity Analysis software (NanoTemper Technologies).

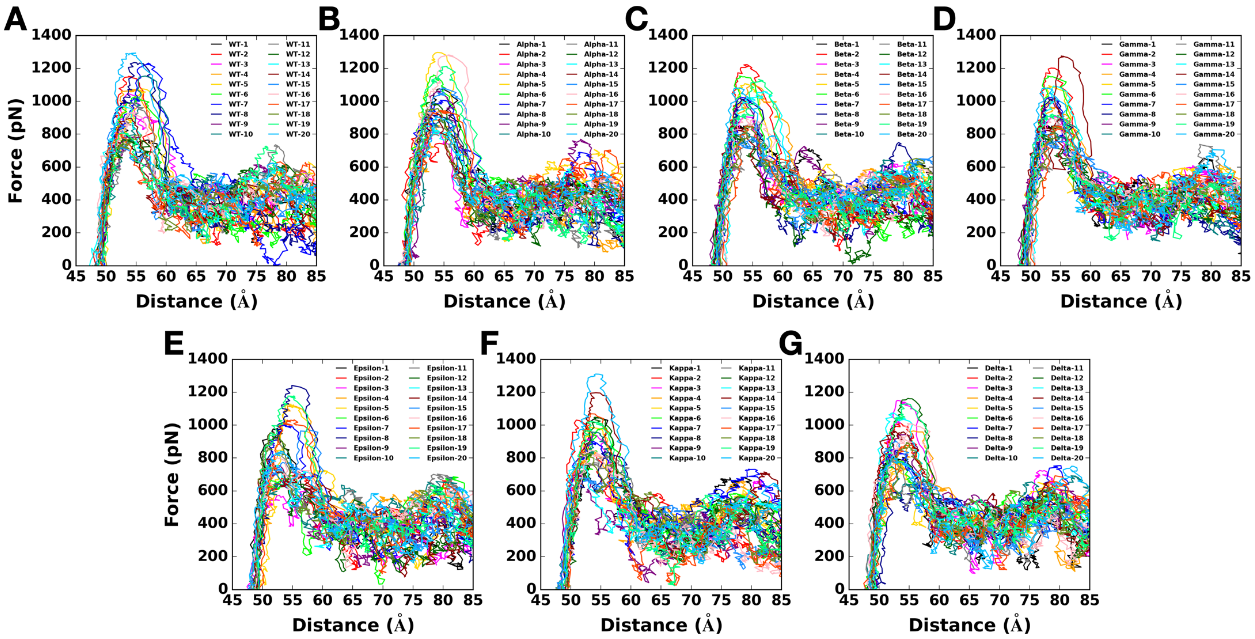


**Figure S1.** Force profiles of 20 replicas of (A) WT, (B) Alpha, (C) Beta, (D) Gamma, (E) Epsilon, (F) Kappa, and (G) Delta as a function of the distance between the COMs of RBD and ACE2.

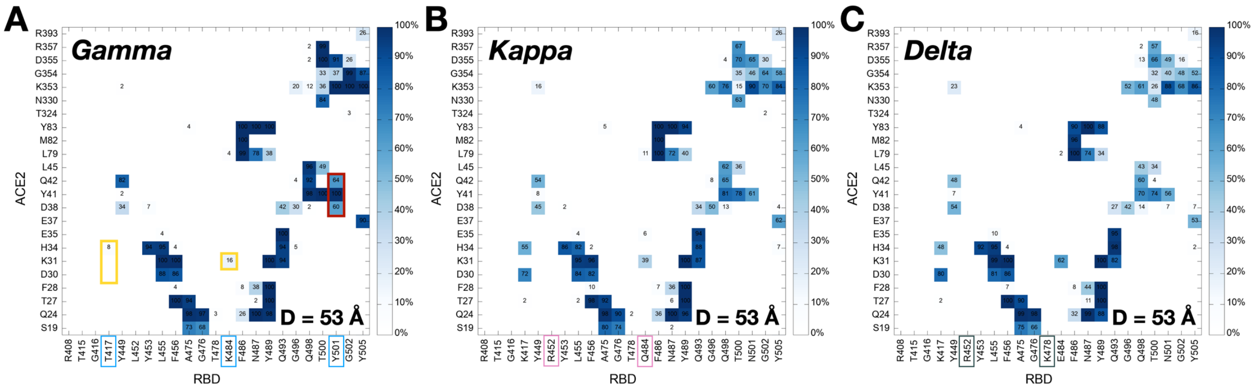


**Figure S2.** Two-dimensional contact maps at $D$ = 53 Å. Mutated RBD residues are labeled as colored boxes: sky blue for Gamma, pink for Kappa, and gray for Delta. The contact frequency is numbered with colors from light blue to dark blue. Dark red and yellow colors on the map respectively represent increased and decreased interactions between RBD and ACE2 upon mutations.

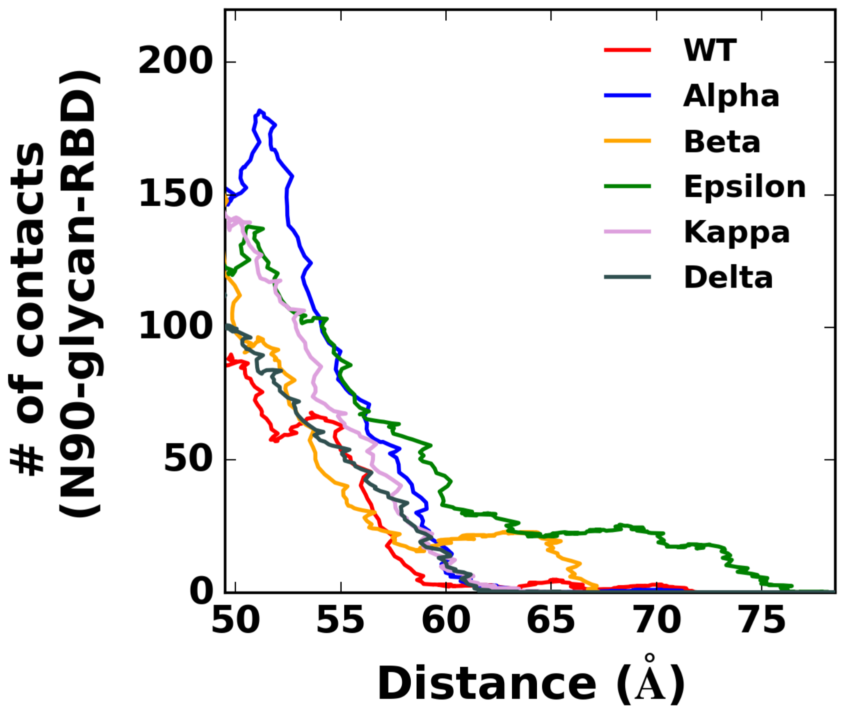


**Figure S3.** The number of heavy atom contacts between RBD and ACE2 N90-glycan. The numbers are counted if any heavy atom of N90-glycan is within 4.5 Å of any residue in RBD. The color scheme is the same as in **Figure 1A**.

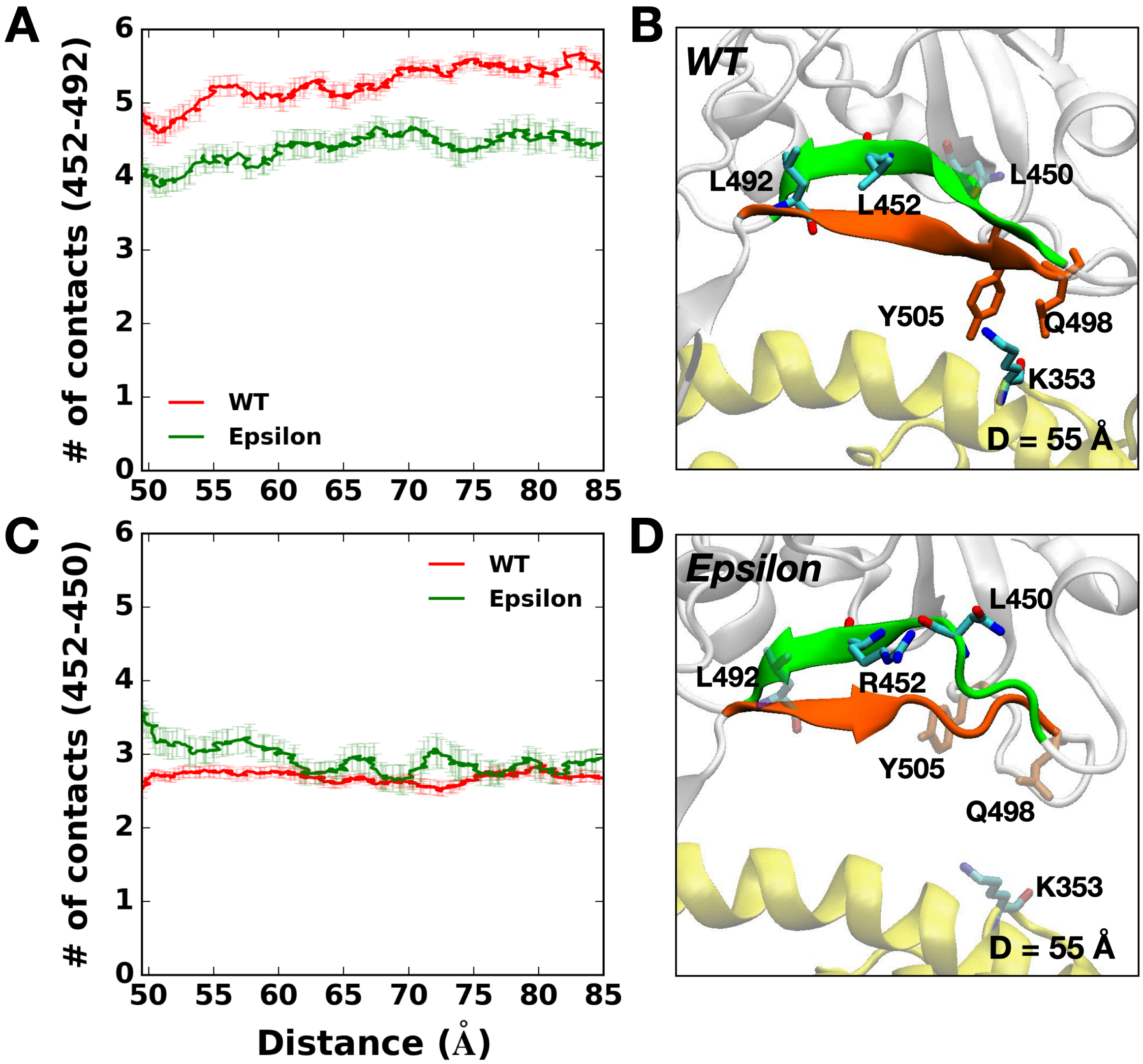


**Figure S4.** The number of heavy atom contacts (A) between RBD residues 452 and 492 and (C) between RBD residues 452 and 450. WT contains L452, and it is mutated to R452 in Epsilon variant. (B) L452 in one β-sheet colored in green interacts with L492 in another β-sheet presented by orange, and such interactions maintain stable secondary structures. (D) R452 of Epsilon variant located in a β-strand colored in green tends to interact with L450 in the same β-strand, having less stable secondary RBD structures by shortening the β-strands. Residues interacting with each other are represented by solid sticks, and those that lost their interaction are shown as transparent sticks.

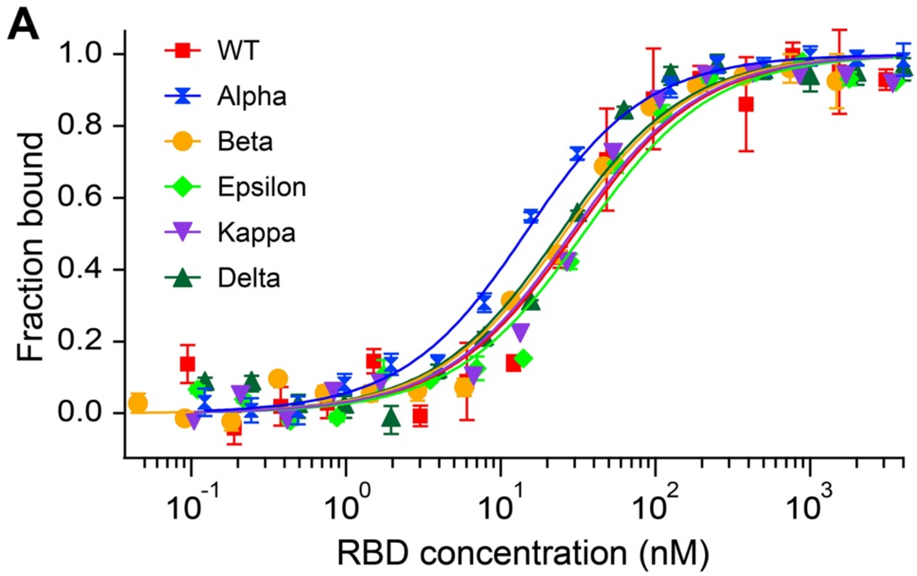


**B**

| RBD variants | $K_d$ (nM) | Standard deviation (± nM) |
|---|---|---|
| WT | 27.5 | 4.8 |
| Alpha | 11.8 | 0.8 |
| Beta | 23.1 | 2.4 |
| Epsilon | 31.7 | 3.9 |
| Kappa | 26.0 | 3.2 |
| Delta | 21.5 | 2.9 |

**Figure S5.** (A) Microscale thermophoresis (MST) analysis of the interaction between ACE2 and six different RBD variants. Error bars represent standard deviations from three to six individual repeat measurements. The binding affinities were determined by fitting the data with the '$K_d$' model of the MO Affinity software. (B) Affinities of ACE2 binding to RBD variants detected by MST. The MST responses were fitted to the 1:1 binding model. The $K_d$ rates are shown as fit ± one standard deviation.

### REFERENCES


1. Woo, H.; Park, S.-J.; Choi, Y. K.; Park, T.; Tanveer, M.; Cao, Y.; Kern, N. R.; Lee, J.; Yeom, M. S.; Croll, T. I.; Seok, C.; Im, W. Developing a Fully Glycosylated Full-Length SARS-CoV-2 Spike Protein Model in a Viral Membrane. *J. Phys. Chem. B* **2020**, *124* (33), 7128-7137.
2. Jo, S.; Kim, T.; Iyer, V. G.; Im, W. CHARMM-GUI: a web-based graphical user interface for CHARMM. *J. Comput. Chem.* **2008**, *29* (11), 1859-1865.
3. Lee, J.; Cheng, X.; Swails, J. M.; Yeom, M. S.; Eastman, P. K.; Lemkul, J. A.; Wei, S.; Buckner, J.; Jeong, J. C.; Qi, Y.; Jo, S.; Pande, V. S.; Case, D. A.; Brooks, C. L.; MacKerell, A. D.; Klauda, J. B.; Im, W. CHARMM-GUI Input Generator for NAMD, GROMACS, AMBER, OpenMM, and CHARMM/OpenMM Simulations Using the CHARMM36 Additive Force Field. *J. Chem. Theory Comput.* **2016**, *12* (1), 405-413.
4. Hatcher, E.; Guvench, O.; MacKerell, A. D. CHARMM Additive All-Atom Force Field for Aldopentofuranoses, Methyl-aldopentofuranosides, and Fructofuranose. *J. Phys. Chem. B* **2009**, *113* (37), 12466-12476.
5. Huang, J.; Rauscher, S.; Nawrocki, G.; Ran, T.; Feig, M.; de Groot, B. L.; Grubmüller, H.; MacKerell, A. D. CHARMM36m: an improved force field for folded and intrinsically disordered proteins. *Nat. Methods* **2017**, *14* (1), 71-73.
6. Jorgensen, W. L.; Chandrasekhar, J.; Madura, J. D.; Impey, R. W.; Klein, M. L. Comparison of simple potential functions for simulating liquid water. *J. Chem. Phys.* **1983**, *79* (2), 926-935.
7. Cao, W.; Dong, C.; Kim, S.; Hou, D.; Tai, W.; Du, L.; Im, W.; Zhang, X. F. Biomechanical characterization of SARS-CoV-2 spike RBD and human ACE2 protein-protein interaction. *Biophys. J.* **2021**, *120* (6), 1011-1019.
8. Phillips, J. C.; Braun, R.; Wang, W.; Gumbart, J.; Tajkhorshid, E.; Villa, E.; Chipot, C.; Skeel, R. D.; Kale, L.; Schulten, K. Scalable molecular dynamics with NAMD. *J. Comput. Chem.* **2005**, *26* (16), 1781-1802.
9. Dion, M.; Rydberg, H.; Schröder, E.; Langreth, D. C.; Lundqvist, B. I. Van der Waals density functional for general geometries. *Phys. Rev. Lett.* **2004**, *92*, 246401.
10. Essmann, U.; Perera, L.; Berkowitz, M. L.; Darden, T.; Lee, H.; Pedersen, L. G. A smooth particle mesh Ewald method. *J. Chem. Phys.* **1995**, *103* (19), 8577-8593.
11. Ryckaert, J.-P.; Ciccotti, G.; Berendsen, H. J. Numerical integration of the cartesian equations of motion of a system with constraints: molecular dynamics of n-alkanes. *J. Comput. Phys.* **1977**, *23* (3), 327-341.
12. Hopkins, C. W.; Le Grand, S.; Walker, R. C.; Roitberg, A. E. Long-time-step molecular dynamics through hydrogen mass repartitioning. *J. Chem. Theory Comput.* **2015**, *11* (4), 1864-1874.
13. Gao, Y.; Lee, J.; Smith, I. P. S.; Lee, H.; Kim, S.; Qi, Y.; Klauda, J. B.; Widmalm, G. r.; Khalid, S.; Im, W. CHARMM-GUI Supports Hydrogen Mass Repartitioning and Different Protonation States of Phosphates in Lipopolysaccharides. *J. Chem. Inf. Model.* **2021**, *61* (2), 831-839.
14. Feller, S. E.; Zhang, Y.; Pastor, R. W.; Brooks, B. R. Constant pressure molecular dynamics simulation: the Langevin piston method. *J. Chem. Phys.* **1995**, *103* (11), 4613-4621.
